# SearcHPV: a novel approach to identify and assemble human papillomavirus-host genomic integration events in cancer

**DOI:** 10.1101/2021.02.11.430847

**Authors:** Lisa M. Pinatti, Wenjin Gu, Yifan Wang, Ahmed El Hossiny, Apurva D. Bhangale, Collin V. Brummel, Thomas E. Carey, Ryan E. Mills, J. Chad Brenner

## Abstract

**Background:** Human papillomavirus (HPV) is a well-established driver of malignant transformation in a number of sites including head and neck, cervical, vulvar, anorectal and penile squamous cell carcinomas; however, the impact of HPV integration into the host human genome on this process remains largely unresolved. This is due to the technical challenge of identifying HPV integration sites, which includes limitations of existing informatics approaches to discover viral-host breakpoints from low read coverage sequencing data.

**Methods:** To overcome this limitation, we developed a new HPV detection pipeline called SearcHPV based on targeted capture technology and applied the algorithm to targeted capture data. We performed an integrated analysis of SearcHPV-defined breakpoints with genome-wide linked read sequencing to identify potential HPV-related structural variations.

**Results:** Through analysis of HPV+ models, we show that SearcHPV detects HPV-host integration sites with a higher sensitivity and specificity than two other commonly used HPV detection callers. SearcHPV uncovered HPV integration sites adjacent to known cancer-related genes including *TP63* and *MYC*, as well as near regions of large structural variation. We further validated the junction contig assembly feature of SearcHPV, which helped to accurately identify viral-host junction breakpoint sequences. We found that viral integration occurred through a variety of DNA repair mechanisms including non-homologous end joining, alternative end joining and microhomology mediated repair.

**Conclusions:** In summary, we show that SearcHPV is a new optimized tool for the accurate detection of HPV-human integration sites from targeted capture DNA sequencing data.

## INTRODUCTION

Human papillomavirus (HPV) is a well-established driver of malignant transformation in a number of cancers, including head and neck squamous cell carcinomas (HNSCC). Although HPV genomic integration is not a normal event in the lifecycle of HPV, it is frequently reported in HPV+ cancers^1–4^ and it may be a contributor to oncogenesis. In cervical cancer, HPV integration increases in incidence during progression from stages of cervical intraepithelial neoplasia (CIN) I/II, CIN III and invasive cancer development.^5^ This process has a variety of impacts on both the HPV and cellular genomes, including disruption of the transcriptional repressor of the HPV oncoproteins E2, leading to increase in genetic instability.^6^ HPV integration occurs within/near cellular genes more often than expected by chance^7^ and has been reported to be associated with structural variations^8^. Recent studies in HNSCCs have also suggested that additional oncogenic mechanisms of HPV integration may exist through direct effects on cancer-related gene expression and generation of hybrid viral-host fusion transcripts.^9^

A wide array of methods has been previously used for the detection of HPV integration. Polymerase chain reaction (PCR)-based methods, such as Detection of Integrated Papillomavirus Sequences PCR (DIPS-PCR)^10^ and Amplification of Papillomavirus Oncogene Transcripts (APOT)^11^, are low sensitivity assays and are limited in their ability to detect the broad spectrum of genomic changes resulting from this process. Next-generation sequencing (NGS) technologies overcomes these limitations. Previous groups have assessed HPV integration within HNSCC tumors in The Cancer Genome Atlas (TCGA) and cell lines by whole-genome sequencing (WGS).^2, 3, 8^ There are a variety of viral integration detection tools developed for WGS data, such as VirusFinder2^12, 13^ and VirusSeq^14^. However, these strategies are designed for a broad range of virus types and require whole genomes to be sequenced at uniform coverage, which can result in a lower sensitivity of detection for specific types of rare viral integration events.

To overcome this issue, others have begun to use HPV targeted capture sequencing.^5, 15–18^ This strategy allows for better coverage of integration sites than an untargeted approach like WGS but requires sensitive and accurate viral-human fusion detection bioinformatic tools, of which the field has been lacking. In our lab, we have found the previously available viral integration callers to have a relatively low validation rate and limitations on the structural information surrounding the fusion sites, which impairs mechanistic studies. Therefore, we set out to generate a novel pipeline specifically for targeted capture sequencing data to serve as a new gold standard in the field.

## MATERIALS AND METHODS

### Targeted Capture Sequencing

DNA from UM-SCC-47 and PDX-294R were submitted to the University of Michigan Advanced Genomics Core for targeted capture sequencing. Targeted capture was performed using a custom designed probe panel with high density coverage of the HPV16 genome, the HPV18/33/35 L2/L1 regions, and over 200 HNSCC-related genes, which are detailed in Heft Neal et. al 2020.^19^ Following library preparation and capture, the samples were sequenced on an Illumina NovaSEQ6000 or HiSEQ4000, respectively, with 300nt paired end run. Data was de-multiplexed and FastQ files were generated.

### Novel Integration Caller (SearcHPV)

The pipeline of SearcHPV has four main steps which are detailed below: (1) Alignment; (2) Genome fusion point calling; (3) Assembly; (4) HPV fusion point calling **(Figure 1)**. The package is available on Github: https://github.com/mills-lab/SearcHPV.

**Figure 1:**
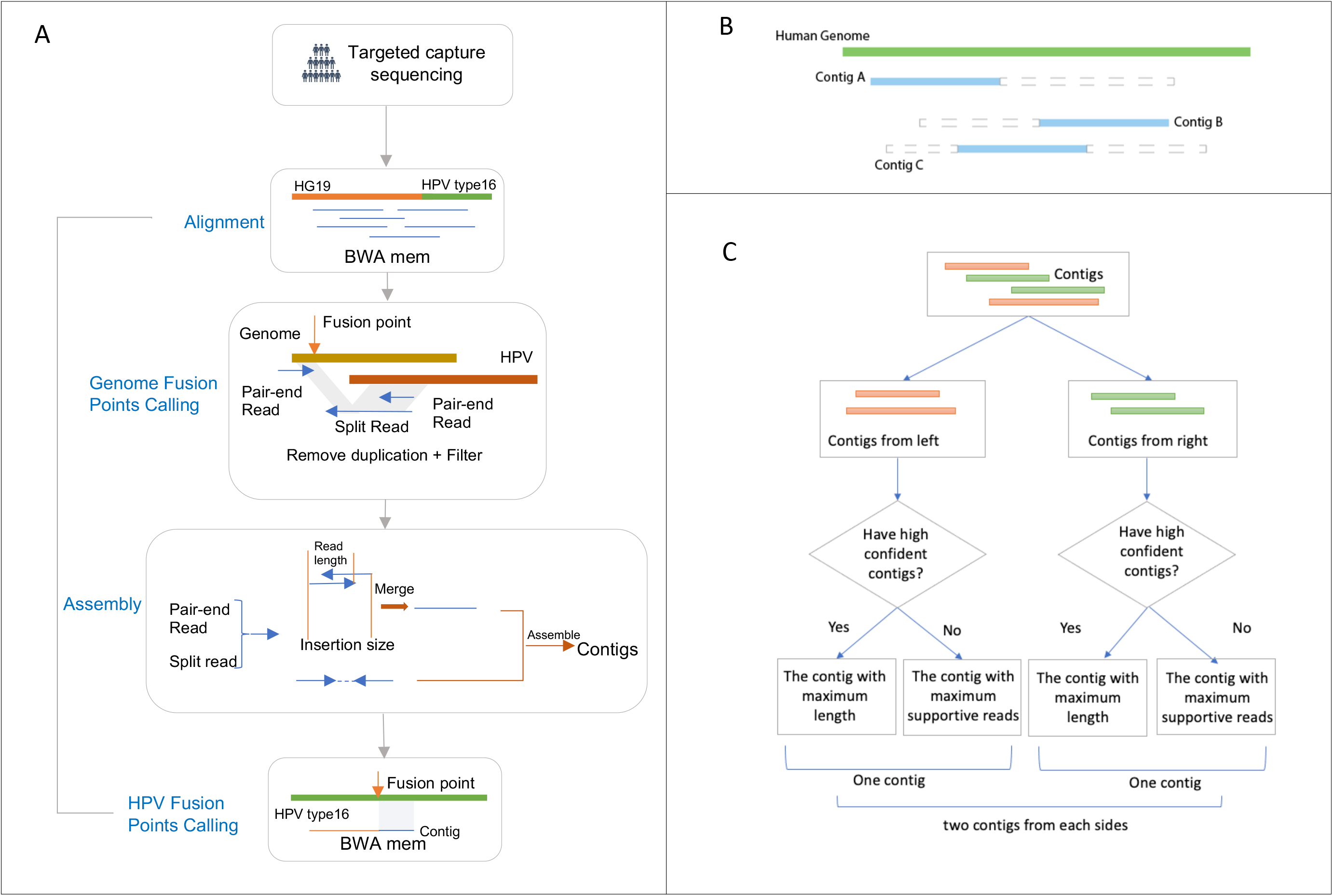
Workflow of SearcHPV. (A) Paired-end reads from targeted capture sequencing were aligned to a catenated Human-HPV reference genome. After removing duplication and filter, fusion points were identified by split reads and pair-end reads. Informative reads were extracted for local assembly. Reads pairs that have overlaps were merged first before assembly. Assembled contigs were aligned to HPV genome to identify the breakpoints on HPV. (B) Contigs were divided to two classes. Blue solid triangle demonstrates the matched region of the contig. Grey dashed triangle demonstrates the clipped region of the contig. Contig A would be assigned to left group and Contig B would be assigned to right group. Contig C would be randomly assigned to left or right group. (C) Workflow for the contig selection procedures for fusion point with multiple candidates contigs. For each fusion point. we report at least one contig and at most two contigs representing two directions.

#### Alignment

The customized reference genome used for alignment was constructed by catenating the HPV16 genome (from Papillomavirus Episteme (PAVE) database^20, 21^) and the human genome reference (1000 Genomes Reference Genome Sequence, hs37d5). We aligned paired-end reads from targeted capture sequencing against the customized reference genome using BWA mem aligner.^22^ Then we performed an indel realignment by Picard Tools^23^ and GATK^24^. Duplications were marked by Picard MarkDuplicates Tool^23^ for the filtering in downstream steps.

#### Genome Fusion Points Calling

To identify the fusion points, we extracted reads with regions matched to HPV16 and filtered those reads to meet these criteria: (1) not secondary alignment; (2) mapping quality greater or equal than 50; (3) not duplicated. Genome fusion points were called by split reads (reads spanning both the human and HPV genomes) and the paired-end reads (reads with one end matched to HPV and the other matched the human genome) at the surrounding region (+/−300bp) (**Figure 1A**). The cut-off criteria for identifying the fusion points were based on empirical practice. We then clustered the integration sites within 100bp to avoid duplicated counting of integration events due to the stochastic nature of read mapping and structural variations.

#### Assembly

To construct longer sequence contigs from individual reals, we extracted supporting split reads and paired-end for local assembly from each integration event. Due to the library preparation methods we implemented for the targeted capture approach, some reads exhibited an insertion size less than 2 x read length, resulting in overlapping read segments. For such events, we first merged these reads using PEAR^25^ and then combined them with other individual reads to perform a local assembly by CAP3^26^ (**Figure 1**).

#### HPV Fusion Point Calling

For each integration event, the assembly algorithm was able to report multiple contigs. We developed a procedure to evaluate and select contigs for each integration event to call HPV fusion point more precisely. First, we aligned the contigs against the human genome and HPV genome separately by BWA mem. If the contig met the following criteria, we marked it as high confidence:

(1) Has at least 10 supportive reads
(2)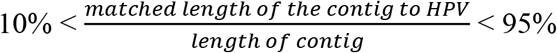

Then we separated the contigs we assembled into two classes: from left side (Contig A in **Fig 1B**) and from right side (Contig B in **Fig 1B**). For each class, if there were high confidence contigs in the class, we selected the contig with maximum length among them, otherwise we selected the contig with most supportive reads. For each insertion event, we reported one contig if it only had contigs from one side and we reported two contigs if it had contigs from both sides (**Figure 1C**). Finally, we identified the fusion points within HPV based on the alignment results of the selected contigs against the HPV genome. The bam/sam file processing in this pipeline was done by Samtools^22^ and the analysis was performed with R 3.6.1^27^ and Python.^28^

## RESULTS

### SearcHPV pipeline

To overcome the limitations of viral integration detection in WGS of detecting rare events, we performed HPV targeted capture sequencing which allows for deeper investigation of these events. Current bioinformatics pipelines available are not designed for this type of data so we developed a novel HPV integration detection tool for targeted capture sequencing data, which we termed “SearcHPV”. Two HPV16+ HNSCC models, UM-SCC-47 and PDX-294R, were subjected to targeted-capture based Illumina sequencing using a custom panel of probes spanning the entire HPV16 genome. The paired end reads then went through the four steps of analysis of SearcHPV: alignment to custom reference genome, genome fusion points calling, local assembly and precise fusion point calling **(Figure 1)**. Analysis of the integration sites in the models using our pipeline SearcHPV showed a high frequency of HPV16 integration with a total of six events in UM-SCC-47 and ninety-eight in PDX-294R (**Figure 2, Table S1-S2**).

**Figure 2:**
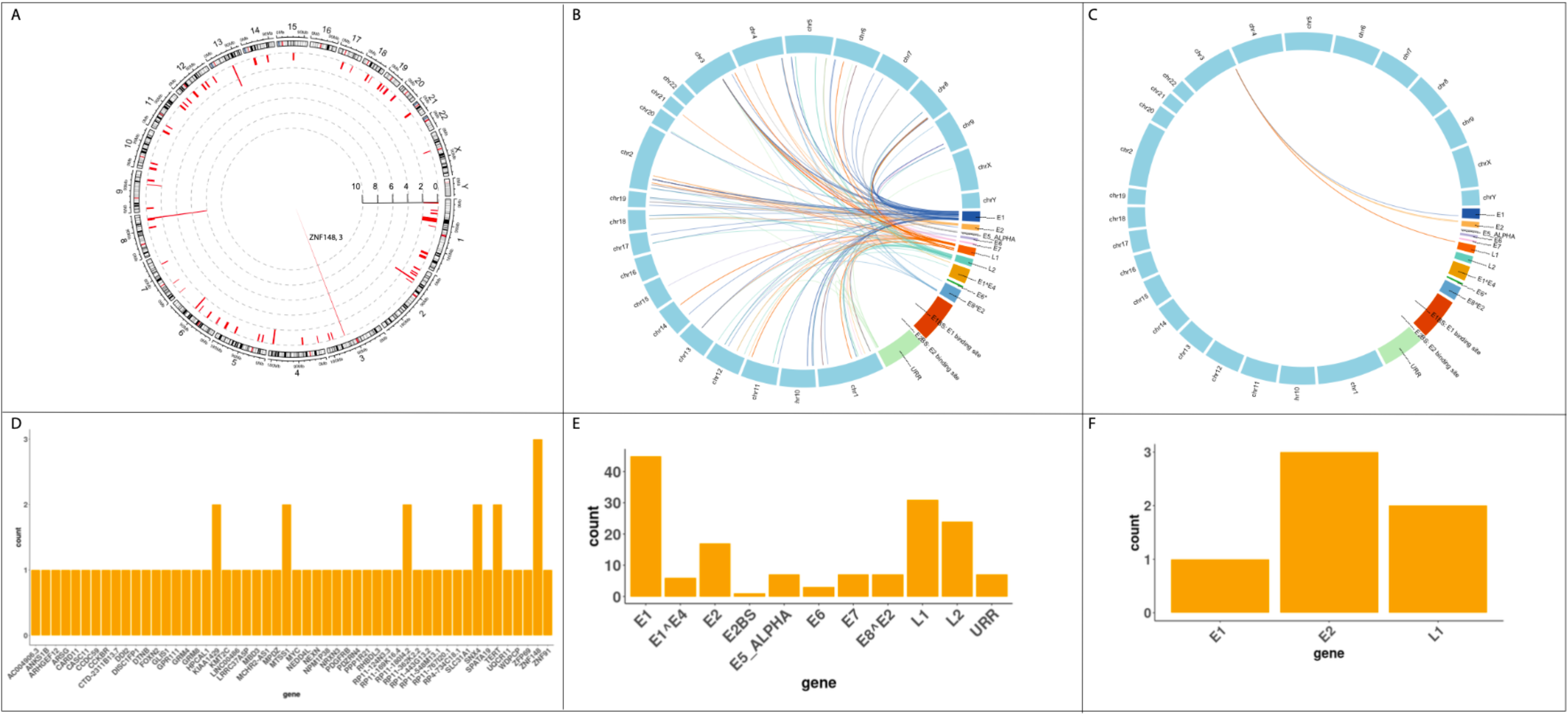
Distribution of breakpoints in the human and HPV genomes called by SearcHPV. (A) Distribution of integration sites in the human genome for PDX-294R. Each bar denotes the count of breakpoints within the region. (B) Links of breakpoints in the human and HPV16 genomes for PDX-294R. (C) Links of breakpoints in the human and HPV16 genomes for UM-SCC-47. (D) Quantification of breakpoint calls in human genes for PDX-294R. (E) Quantification of breakpoints calls in the HPV16 genes for PDX-294R. (F) Quantification of breakpoint calls in the HPV16 genes for UM-SCC-47.

### Comparison to other integration callers and confirmation of integration sites

In addition to using SearcHPV, we used two previously developed integration callers, VirusFinder2 and VirusSeq to independently call integration events in both UM-SCC-47 and PDX-294R (**Figure 3, Tables S3-4**). We found that SearcHPV called HPV integration events at a much higher rate than either previous caller. There were a large number of sites that were only identified by SearcHPV (n=76). In order to assess the accuracy of each caller, we performed PCR on source genomic DNA followed by Sanger sequencing with primers spanning the HPV-human junction sites predicted by the callers (**Figure 3C. S1, Table S5**). We tested all integration sites with sufficient sequence complexity for primer design (n=46), twenty-five of which were unique to SearcHPV and five which were unique to VirusSeq. VirusFinder2 does not allow for local assembly of the integration junctions which rendered us unable to test these sites. Sites unique to SearcHPV had a confirmation rate of 18/25 (72%). The confirmation rate of high confidence SearcHPV sites was higher than that for low confidence sites (23/31 (74%) versus 4/7 (57%)). In contrast, only 2/5 (40%) sites unique to VirusSeq could be confirmed.

**Figure 3:**
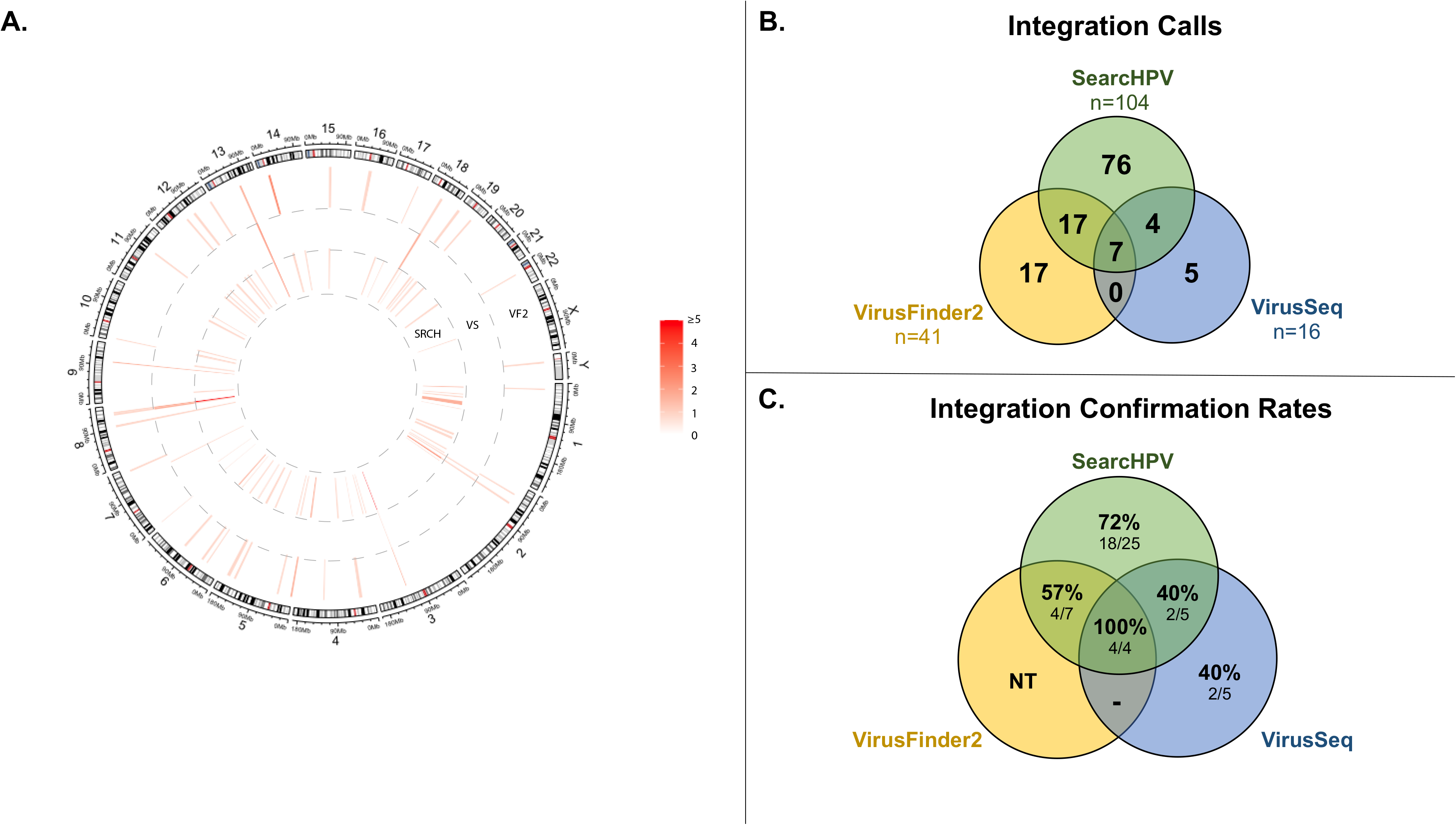
Comparison of integration sites called by SearcHPV, VirusSeq and VirusFinder2 in both models. (A) Each bar denotes an integration site. The colormap shows the count of the integration sites. (B) Number of integration sites called by each program. (C) PCR confirmation rate of sites called by each program.

### Localization of integration sites

We next examined the integration sites detected by SearcHPV. The six integration sites discovered in UM-SCC-47 were clustered on chromosome 3q28 within/near the cellular gene *TP63* and either involved the HPV16 genes E1, E2 or L1. The integration sites fell within intron 10, intron 12 and exon 14. One additional integration site was 8.6 kb downstream of the *TP63* coding region.

Within PDX-294R, HPV16 integration sites were identified across 21 different chromosomes, occurring most frequently on chromosome 3. For the 98 integration events of PDX-294R, we identified 142 breakpoints in the HPV genome. The most frequently involved HPV genes were E1 (45/142 (32%)) and L1 (31/142 (22%)). Most of the integration sites mapped to within/near (<50 kb) a known cellular gene (89/98 (91%)). Of the sites that fell within a gene, the majority of integrations took place within an intronic region (33/42 (78%)). Although the integration sites were scattered throughout the human genome, we saw examples of closely clustered sites around cancer-relevant genes, including *ZNF148* and *SNX4* on chromosome 3q21.2, *MYC* on chromosome 8q24.21 and *FOXN2* on chromosome 2p16.3.

### Association of integration sites and large-scale duplications

We predicted that the complex integration sites we discovered in UM-SCC-47 and PDX-294R would be associated with large-scale structural alterations of the genome, such as rearrangements, deletions and duplications. To identify these alterations, we subjected UM-SCC-47 and PDX-294R to 10X linked-read sequencing. We generated over 1 billion reads for each sample (**Table S6**), with phase blocks (contiguous blocks of DNA from the same allele) of up to 28.9M and 3.8M bases in length for UM-SCC-47 and PDX-294R, respectively (**Figure S2**). This led to the identification of 444 high confidence large structural events in UM-SCC-47 and 126 events in the PDX-294R model. We then performed integrated analysis with our SearcHPV results. There was a 130 kb duplication surrounding the integration events in *TP63* in UM-SCC-47 (**Figure 4A**). In PDX-294R, 38/98 (39%) integration sites were within a region that contained a large-scale duplication, while the other 50 integration events fell outside regions of large structural variation. This suggested that in this PDX model, 38/126 (30%) large structural events were potentially induced during HPV integration. For example, the clusters of integration events surrounding *ZNF148* and *SNX4*, *MYC*, as well as *FOXN2* were also associated with large genomic duplications (**Figure 4B-C**).

**Figure 4:**
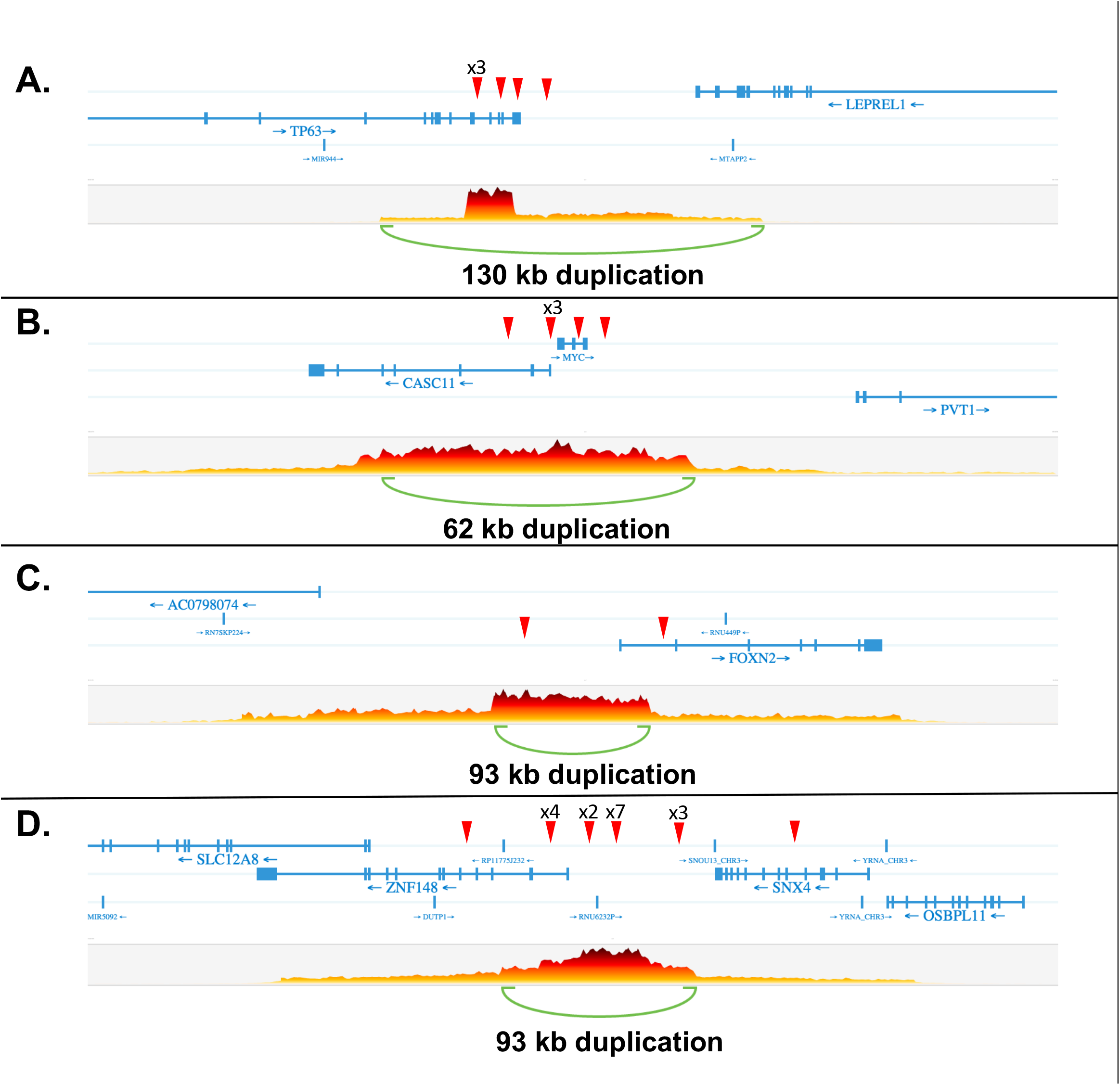
Genomic duplications associated with HPV integration in UM-SCC-47 (A) and PDX-294R (B-D). Red arrows indicate integration site. Each plot shows the number of overlapping barcodes observed in sequencing reads of that region.

### Microhomology at junction sites

Finally, to evaluate possible mechanisms of DNA repair-mediated integration, we examined the degree of sequence overlap between the genomes at each junction sites that covered by contigs. We saw three types of junction points: those with a gap of unmapped sequence between the human and HPV genomes, those that had a clean breakpoint between the genomes, and those with sequence that could be mapped to both genomes (**Figure 5A**). The majority of junction sites in both samples had at least some degree of microhomology (58%) (**Figure 5B-C**). Integration sites with clean breaks (0 bp overlap) and 3 bp of overlap were the most frequently seen junctions in PDX-294R, but there was a wide range of levels seen. There was also a large number of junctions with gaps between the human and HPV genomes ranging from 1 - 54 bp long.

**Figure 5:**
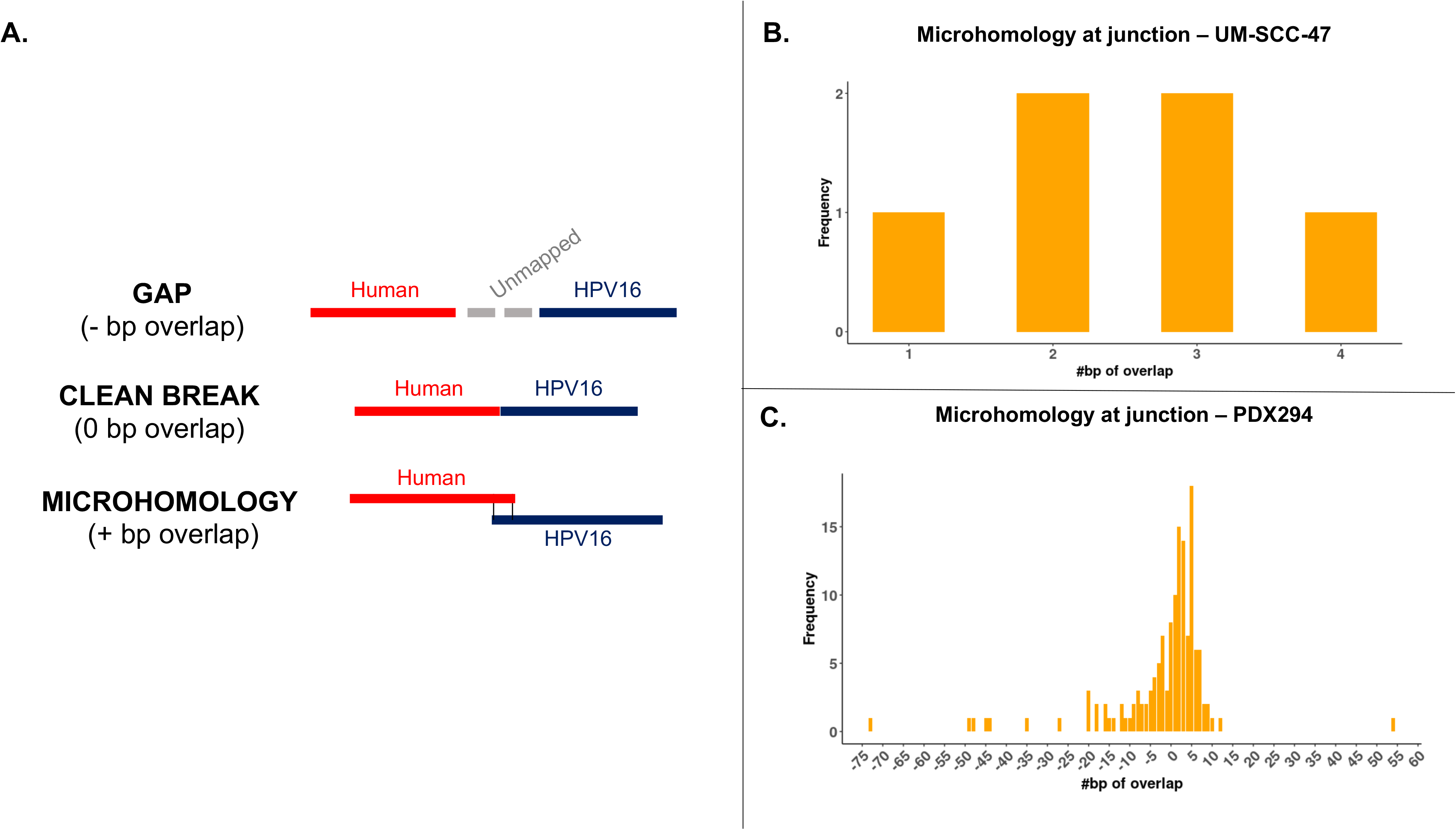
Microhomology at junction points. (A) The three types of junction points. (B) Level of microhomology (in bp) in UM-SCC-47. (C) Level of microhomology (in bp) in PDX-294R. Junctions with a gap are shown as negative numbers.

## Discussion

We developed a novel bioinformatics pipeline that we termed “SearcHPV” and show that it operated in a more accurate and efficient manner than existing pipelines on targeted capture sequencing data. The software also has the advantage of performing local contig assembly around the junction sites, which simplifies downstream confirmation experiments. We used our new caller to interrogate the integration sites found in two HNSCC models in order to compare the accuracy of our caller to the existing pipelines. We then evaluated the genomic effects of these integrations on a larger scale by 10X linked-reads sequencing to identify the role of HPV integration in driving structural variation in the tumor genome.

Using SearcHPV, we were able to investigate the HPV-human integration events present in UM-SCC-47 and PDX-294R. Importantly, UM-SCC-47 has been previously assessed for HPV integration by a variety of methods^8, 29–32^, which we leveraged as ground truth knowledge to validate our integration caller. All previous studies were in agreement that HPV16 is integrated within the cellular gene *TP63*, although the exact number of sites and locations within the gene varied by study. In this study, SearcHPV also called HPV integration sites within *TP63*. We found integrations of E1, E2 and L1 within *TP63* intron 10, L1 within intron 12 and E2 within *TP63* exon 14. These integration sites were also detected using DIPS-PCR^32^ and/or WGS^8^ with the exception of E1 into intron 10, which was unique to our caller and confirmed by direct PCR. It is possible that the integration sites detected in this sample represent multiple fragments of one larger integration site. There were additional sites called by other WGS studies that we did not detect (intron 9^8^ and exon 7^31^), although it is possible that alternate clonal populations grew out due to different selective pressures in different laboratories. Nonetheless, the analysis clearly demonstrated that SearcHPV was able to detect a well-established HPV insertion site.

In contrast to UM-SCC-47, to our knowledge PDX-294R has not been previously analyzed for viral-host integration sites and therefore represented a true discovery case. We identified widespread HPV integration sites throughout the host genome and also observed that 66% of integration sites were found within or near genes. This aligns with previous reports that integrations are detected in host genes more frequently than expected by chance.^2, 3, 7, 33^ One particularly interesting cluster of integration events surrounded the cellular proto-oncogene *MYC*. Importantly, *MYC* has been identified as a potential hotspot for HPV integration^7, 34^ and the junctions we detected in/near this gene had 2-4 bp of microhomology, potentially driving this observation. Accordingly, an HPV-integration related promoter duplication event, which may be expected to drive expression, would be consistent with a novel genetic mechanism to drive expression of this oncogene.

*TP63* has also been reported to be a hotspot for HPV integration, as it has been recorded in multiple samples besides UM-SCC-47.^3, 7, 35, 36^ There is a high degree of microhomology between HPV16 and this gene. Given the high frequency of molecular alterations in the epidermal differentiation pathway (e.g. *NOTCH1/2*, *TP63* and *ZNF750*) in HPV+ HNSCCs, this data supports HPV integration as a pivotal mechanism of viral-driven oncogenesis in this model.^37^

HPV integration sites have been associated with structural variations in the human genome^3, 8, 37^, which supports an additional genetic mechanism as to why HPV integration sites may often be detected adjacent to host cancer-related genes. These structural variation events are thought to be due to the rolling circle amplification that takes place at the integration breakpoint, leading to the formation of amplified segments of genomic sequence flanked by HPV segments.^8, 38^ Our data are consistent with these previous reports in that approximately half of the integration events we discovered were associated with a large-scale amplification. It is unclear why only some integration sites were associated with structural variants, but it is possible that an alternative mechanism of integration occurred.^38^

Importantly, this observation that HPV integration events tended to be enriched in cellular genes could result from multiple different mechanisms. Integration could occur preferentially in regions of open chromatin during cell replication and keratinocyte differentiation. Other potential mechanisms are: 1) that HPV integration is directed to specific host genes by homology, or 2) that HPV integration is random, but events that are advantageous for oncogenesis are clonally selected and expanded, implicating non-homology based DNA repair mechanisms. Therefore, to help resolve differences in the mechanism of integration, we assessed microhomology at the HPV-human junction points. The majority of breakpoints had some level of microhomology. The most frequent levels of overlap were 0 and 3 bp, which potentially implicates non-homologous end joining (NHEJ) in repair at these sites, since this pathway most frequently results in 0-5 bp of overlap.^39^ There were also a number of junction sites that demonstrated a gap of inserted sequence between the HPV and human genomes. It has been described that during polymerase theta-mediated end joining (TMEJ), stretches of 3-30 bp are frequently inserted at the site of repair, possibly accounting for these sites.^40^ However, given the relatively small number of events we examined, we expect that future analysis with our pipeline will help resolve the specific role of each DNA repair pathway in HPV-human fusion breakpoints.

Overall, our new HPV detection pipeline SearchHPV overcomes a gap in the field of viral-host integration analysis. While the performance of SearcHPV has only been examined on two models, in the future, we expect that the application of this pipeline in large HPV+ cancer tissue cohorts will help advance our understanding of the potential oncogenic mechanisms associated with viral integration. With the emerging set of tools such as SearcHPV, we believe the field is now primed to make major advances in the understanding of HPV-driven pathogenesis, some of which may lead to the development of novel biomarkers and/or treatment paradigms.

## ACKNOWLEDGMENTS

We would like to thank the University of Michigan Advanced Genomics Core for carrying out the targeted capture sequencing and 10X linked read sequencing. We thank Dr. Tom Wilson for discussions of the data.

---

**Figure S1:**
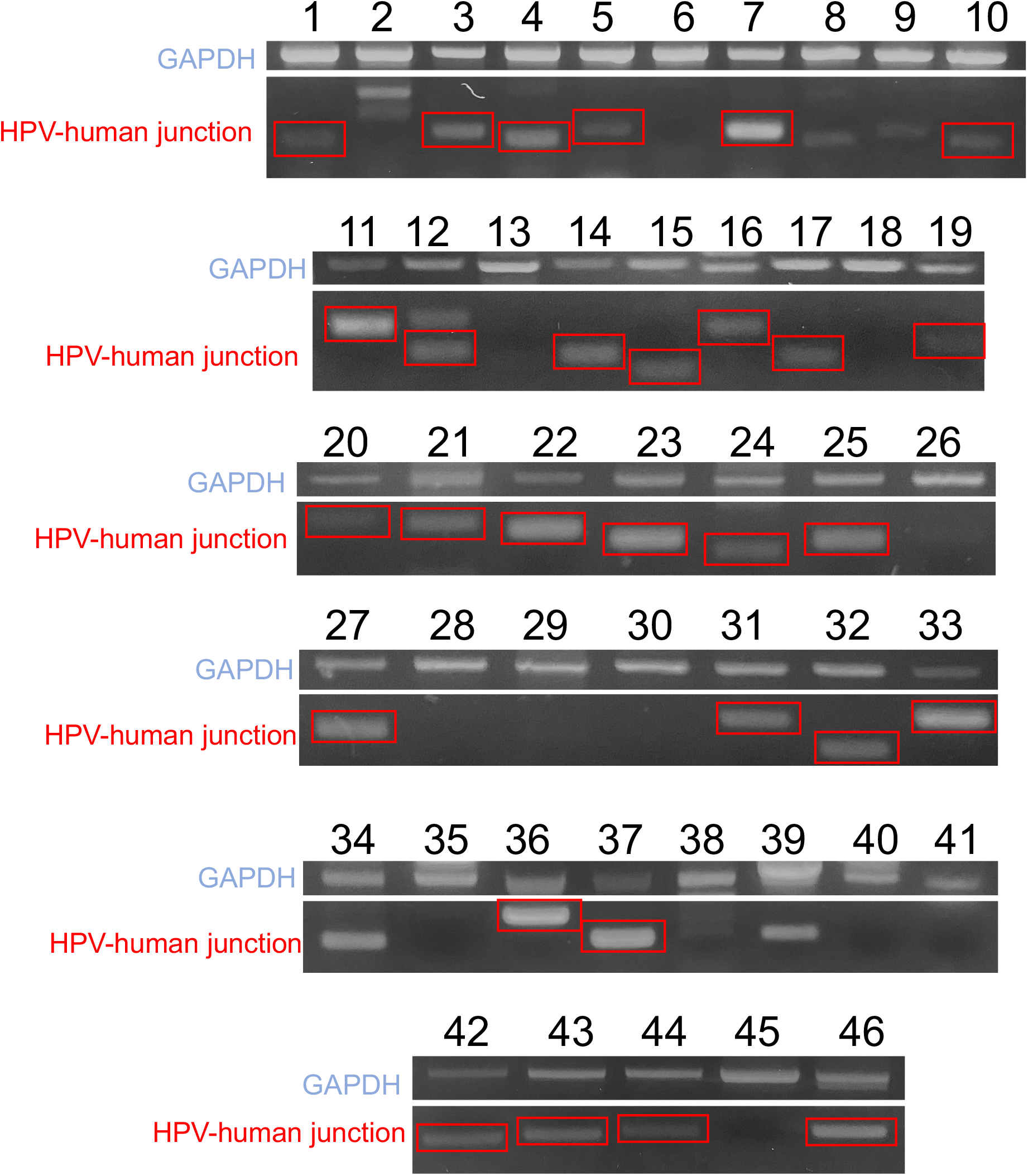
PCR validation gel electrophoresis. Top band of each row shows GAPDH (535 bp), bottom bands represent predicted HPV-human junctions (ranging from 70-250 bp). Red boxes demonstrate bands that appeared at the correct molecular weight and were validated by Sanger sequencing.

**Figure S2:**
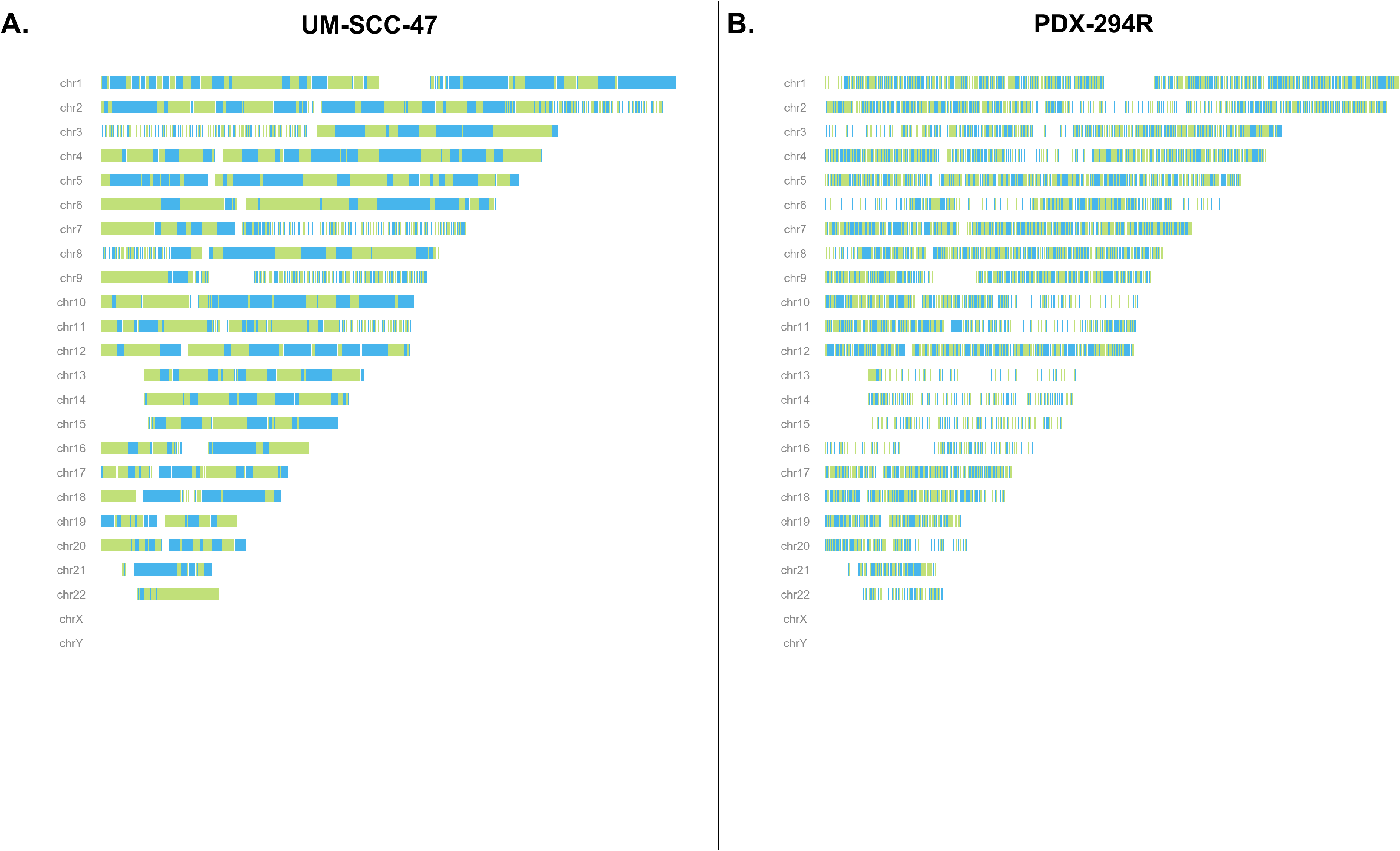
Linked read SNP phase plots for UM-SCC-47 (A) and PDX-294R (B) genomes. Alternating colors represent different phase blocks, which are contiguous blocks of DNA from the same allele based on differential SNP phasing performed by LongRanger software.

