## Supplemental Info for "SearcHPV: a novel approach to identify and assemble human papillomavirus-host genomic integration events in cancer"

**Expanded Materials and Methods:**

**Cell Line Model:** UM-SCC-47 was previously derived in our lab from a surgical resection of a previously untreated p16+ T3N1M0 carcinoma of the lateral tongue in a 53-year-old male smoker.[1] The patient died within a year of diagnosis. Subsequent HPV testing demonstrated the cell line to be p16+ and HPV16+.[2, 3]

**PDX Model:** Flash frozen tissue from an HPV16+ OPSCC PDX model (PDX-932174-294-R, subsequently abbreviated PDX-294R) was obtained from the National Cancer Institute Patient-Derived Models Repository (NCI-PDMR), NCI-Frederick, Frederick National Laboratory for Cancer Research (Frederick, MD) – https://pdmr.cancer.gov/. The PDX was derived from a base of tongue squamous cell carcinoma from a 62-year-old, treatment naive male patient.

**DNA Isolation:** High molecular weight DNA was isolated by treating the samples overnight at 37° with lysis buffer (10 mM Tris-HCl, 400 mM NaCl, 2 mM EDTA), 10% SDS, RNase A and a proteinase K solution (1 mg/mL Proteinase K, 1% SDS, 2 mM EDTA). DNA was then salted out of the solution with 5 M NaCl for 1 hour at 4° and precipitated with ice cold ethanol for 5 hours at -20°C. High molecular weight DNA was eluted in TE buffer; the quality and integrity of the DNA was assessed using the Tapestation Genomic DNA ScreenTape kit (Agilent, Santa Clara, CA).

**Other Integration Callers:**

***VirusSeq***

We installed and ran VirusSeq following the user guide using the default parameter settings. We first installed the MOSAIK aligner.[4] As VirusSeq did not have a feature for users to customize the reference genome, we indexed and used the provided built-in reference database (GIB-V[5], hg19 and hybrid reference genome concatenated by hg19 and 17 viral genomes). VirusSeq aligned the paired end reads to the catenated reference genome by MOSAIK and extracted and clustered split reads using a Perl script (Spanner_cross_converter.pl). Integration sites were detected by another Perl script (VirusSeq_Integration.pl).

***VirusFinder2***

To install VirusFinder2, we first installed all the third-party tools and Perl modules required by VirusFinder2. As required by VirusFinder2, we indexed the human reference genome (hs37d5) by Bowtie2[6] and Blast+[7], as well as the virus database (DB[8]) suggested by VirusFinder2 using Blast+. VirusFinder2 used Bowtie2 to align raw reads against the human reference genome. The informative reads were extracted and assembled to contigs using Trinity.[9] By mapping contigs to the virus reference database, VirusFinder2 detected the virus and identified the virus type. It then applied the VERSE algorithm to customize the reference genome. First, it extracted the viral reads and mapped them to the virus reference genome to modify the variants on the virus reference genome. It next catenated the human reference genome with customized virus reference genome and mapped viral reads to this new reference to extract virus insertion-harboring regions. By mapping raw reads to these regions, VERSE detected the variants in these regions and modified the genomic regions. With the customized reference genome, VirusFinder2 mapped informative reads to the reference genome by BWA[10] and detected the viral integrations and breakpoints by

SVDetect[11] and CREST.[12] The detailed parameter settings for VirusFinder2 can be found in **Table S7**.

**Sanger Sequencing:** Primer sets (n=46) were designed to amplify across the predicted HPV-human junctions from the contigs generated by the integration callers (**Table S8**). Primers were designed using NCBI Primer-BLAST. PCR was performed using each of the 46 primer sets multiplexed with *GAPDH* control primers using 50 ng DNA and Platinum Taq (Thermo Fisher Scientific, Waltham, MA) following the manufacturer's instructions. The PCR products were run by gel electrophoresis on a 1.5% agarose gel, followed by isolation of DNA from the bands at the predicted molecular weights using the Qiaquick Gel Extraction Kit (Qiagen, Hilden, Germany). These products were sent for Sanger sequencing at Eurofins Genomics (Louisville, KY) and mapped back to the predicted sequence to confirm sequence identity.

**Characterization of Integration Calls:** Circos plots detailing integration sites were generated using the Circlize package[13] in R 3.6.1.[14] Distance of each integration site from genes was calculated based on NCBI RefSeq genes (Release 105.20190906).

**10X Linked Reads Sequencing:** High-molecular weight DNA from UM-SCC-47 and PDX-294R was submitted to the University of Michigan Advanced Genomics Core for 10x-based linked read library generation and sequencing on an Illumina NovaSeq6000 with 300nt paired end run. Samples were de-multiplexed and FastQ files with matched index files were generated using Long Ranger Version 2.2.2. Data was visualized using the Loupe software package, Version2.1.1 (2.4). Structural variation calls were considered high confidence if they occurred in unambiguous regions of the reference genome and there were 3 or more supporting sequencing barcodes detected at the site. The raw data was deposited to the sequencing read archives under identification number: PRJNA668771.
